# Compressed timeline of recent experience in monkey lPFC

**DOI:** 10.1101/126219

**Authors:** Zoran Tiganj, Jason A. Cromer, Jefferson E. Roy, Earl K. Miller, Marc W. Howard

**Affiliations:** Center for Memory and Brain, Department of Psychological and Brain Sciences, Boston University, Boston, MA 02215; Department of Brain and Cognitive Sciences, Massachusetts Institute of Technology, Cambridge, MA 02139

## Abstract

Cognitive theories suggest that working memory maintains not only the identity of recently-presented stimuli but also a sense of the elapsed time since the stimuli were presented. Previous studies of the neural underpinnings of working memory have focused on sustained firing, which can account for maintenance of the stimulus identity, but not for representation of the elapsed time. We analyzed single-unit recordings from the lateral prefrontal cortex (lPFC) of two macaque monkeys during performance of a delayed-match-to-category task. Each sample stimulus triggered a consistent sequence of neurons, with each neuron in the sequence firing during a circumscribed period of time. These sequences of neurons encoded both stimulus identity and the elapsed time. The encoding of the elapsed time became less precise as the sample stimulus receded into the past. These findings suggest that working memory includes a compressed timeline of what happened when, consistent with longstanding cognitive theories of human memory.

**Significance Statement:** Place cells in the hippocampus and other brain regions provide basis functions to support the dimension of physical space. Time cells, which activate sequentially provide analogous support for the dimension of time. We observed time cells in the macaque lPFC during a working memory task. The time cells we observed were stimulus specific meaning that they provide not only information about timing, but also conjunctively code what and when information. This representation thus constitutes a manifold with both temporal dimension and a stimulus-coding dimension that could support working memory. These temporal basis functions maintain a logarithmically-compressed timeline of the recent past, providing strong empirical support to long-standing cognitive theories of human memory.

## Introduction

Theories of human memory have long suggested that memory depends on a representation of the recent past in which events are organized on a compressed timeline (James, 1890; Crowder, 1976; Brown, Neath, & Chater, 2007; Howard, Shankar, Aue, & Criss, 2015). This implies that memory provides access to what happened when; a neural representation supporting a timeline should enable reconstruction of the chronological order of previous stimuli as well as their identity. If the timeline is compressed, then as stimuli recede into the past, their time of occurrence and identity is represented with less and less accuracy.

The neural underpinnings of working memory are studied using tasks which require maintenance of a small amount of information across a (typically brief) delay interval. Most neural models assume that working memory maintenance relies on sustained firing of neurons (Goldman-Rakic, 1995; Goldman, 2009; Egorov, Hamam, Fransén, Hasselmo, & Alonso, 2002). According to this model, when a to-be-remembered stimulus is presented, it activates a specific population of neurons that remain firing at an elevated rate for as long as necessary until the information is no longer required. A great deal of work in computational neuroscience has developed mechanisms for sustained stimulus-specific firing at the level of circuits, channels, and LFPs (Amit & Brunel, 1997; Compte, Brunel, Goldman-Rakic, & Wang, 2000; Durstewitz, Seamans, & Sejnowski, 2000; Chaudhuri & Fiete, 2016; Lundqvist, Herman, & Lansner, 2011; Mongillo, Barak, & Tsodyks, 2008; Sandberg, Tegnér, & Lansner, 2003). However, if the firing rate is constant while the stimulus is maintained in working memory, then information about the passage of time is lost. Thus a memory representation based on sustained firing is not sufficient to represent information about time.

Time cells, neurons that fire sequentially, each for a circumscribed period of time, during the delay interval of a memory task (Pastalkova, Itskov, Amarasingham, & Buzsaki, 2008; MacDonald, Lepage, Eden, & Eichenbaum, 2011), provide a neural representation that includes information about time. By examining which time cell is firing at a particular moment, one can reconstruct how far in the past the delay began. Behavioral work on timing shows that the accuracy in estimating the elapsed time decreases with the amount of time to be estimated (e.g., Rakitin et al., 1998; Lewis & Miall, 2009). Two properties of time cells are consistent with an analogous decrease in temporal accuracy. First, time fields later in the sequence should be more broad (i.e., less precise). Second, there should be more neurons with time fields early in the delay and fewer neurons representing times further in the past. Both of these properties have been observed, primarily in rodent work, for time cells in the hippocampus (Howard et al., 2014; Salz et al., 2016), entorhinal cortex (Kraus et al., 2015), medial prefrontal cortex (mPFC) (Tiganj, Kim, Jung, & Howard, 2016), and striatum (Jin, Fujii, & Graybiel, 2009; Mello, Soares, & Paton, 2015; Akhlaghpour et al., 2016).

The cognitive models for a compressed timeline predict that distinct sequences of time cells should be triggered by distinct stimuli. In this way, one could directly read off not only what stimulus occurred in the past, but also how far in the past that stimulus was presented. Despite some dramatic evidence for sequence memory in primate lPFC (Ninokura, Mushi-ake, & Tanji, 2003, 2004), thus far there has been little evidence for conjunctive “what and when” information in populations of time cells. Earlier studies of time cells have not observed stimulus-specific time cell firing (e.g., Akhlaghpour et al., 2016 but see MacDonald, Carrow, Place, & Eichenbaum, 2013) leading some theorists to hypothesize that what and when information are maintained separately (Friston & Buzsáki, 2016) in much the same way that what and where information are presumably segregated in the visual system.

## Methods

This paper reports reanalysis of data initially described in Cromer, Roy, and Miller (2010). More detailed descriptions of the behavioral and recording methods can be found in that paper.

### Behavioral task

Two macaque monkeys, one male and one female, performed a delayed match to category task. The stimuli were chosen from one of two independent category sets; each category set consisted of two categories. One category set was ANIMALS, which consisted of the categories DOGS and CATS. The other category set was CARS, which consisted of the categories SPORTS CARS and SEDANS. Stimuli were constructed as morphed images, composed from a mixture of two prototype images each taken from a different category within the same category set (Figure 2A-B). The test stimulus was always chosen to be from the same category set as the sample stimulus, but it could come from either the same or from a different category within that category set.

Each trial was initiated by the monkey grabbing a response bar. The trials started with a 1000 ms fixation period, during which a white cross was presented in the middle on the screen. The monkey was required to fixate on it. The fixation period was followed by a 600 ms sample stimulus presentations and then by a 1000 ms delay interval during which the monkey had to maintain a memory of the stimulus category in order to be able to successfully complete the task and obtain a reward. The delay period was followed by a test stimulus presentation lasting for another 600 ms.

On match trials, when the test stimulus was from the same category as the sample stimulus, the monkey had to release the response bar within 600 ms of the test stimulus presentation. On nonmatch trials, the monkey had to continue holding the bar during the test stimulus presentation and during a subsequent 600 ms delay interval followed by a second test image. The second test image was always a category match to the sample image (see Figure 2C for a block diagram showing the behavioral protocol). The performance of each monkey was correct in more than 80% of trials.

### Electrophysiological Recordings

Neural recordings were made using up to 16 individual, epoxy-coated tungsten electrodes (FHC Inc., Bowdoin, ME) positioned over the lPFC. Spike sorting was performed on digitalized waveforms using principal components analysis (Offline Sorter, Plexon Inc., Dallas, TX). A total of 500 isolated units was recorded from the two animals. Recordings of 455 out of these 500 units were already used for the analysis published in (Cromer et al., 2010). Eye movements were recorded using an infrared eye tracking system (Iscan, Burlington, MA).

### Identifying temporal and stimulus selectivity using Maximum likelihood

We used a maximum likelihood approach to evaluate whether time and/or stimulus identity were encoded in the firing of the recorded cells. These methods build on analysis methods used to identify time cells in rodent hippocampus and mPFC (Salz et al., 2016; Tiganj et al., 2016). Here we expanded the approach to include the identify of the stimulus in the modeled firing rates. This enables the method to identify conjunctive, stimulus-specific time cells. In each trial we only analyzed the 1600 ms starting from presentation of the sample and terminating at the presentation of the next stimulus. This interval includes the 600 ms presentation of the sample stimulus and a 1000 ms blank delay interval. The spike trains of each cell were fitted with models that included different variables, such as time and stimulus identity. The parameter space of these models was systematically explored in order to compute the maximum likelihood fit. To find the best-fitting model the parameter space was iteratively searched using a combination of particle swarming and the Quasi-Newton method. Particle swarming was performed first (with the swarm size equal to 50) and its output was used to initialize the Quasi-Newton method which was performed second (the number of maximum function evaluations was set to 10000). The computations were implemented in Matlab 2016a. To avoid solutions that converged to a local minimum, the fitting procedure was repeated until the algorithm did not result with better likelihood for at least five consecutive runs. As a preprocessing step, spike trains were downsampled to 1 ms temporal resolution such that if a spike was observed in a particular 1 ms time bin, the corresponding data point was set to 1, otherwise it was set to 0. The maximum likelihood was computed for each recorded cell using all available trials.

### Identifying temporal receptive fields

We first identified cells whose firing was modulated by the passage of time. This was done by comparing the maximum likelihood of the fits from two different models, one containing a Gaussian-shaped time field and the other containing only a constant term. The model with Gaussian-shaped time fields had a set of parameters Θ, which consisted of a constant term *a*_0_, the amplitude of the time fields *a*_1_, the mean *μ*_*t*_ and standard deviation *σ*_*t*_ of the Gaussian time field. With this model the probability of a spike at any given time point *t* was given as:

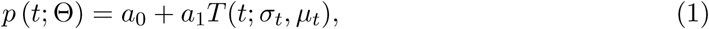

where the Gaussian-shaped time field *T*(*t*;*σ*_*t*_, *μ*_*t*_

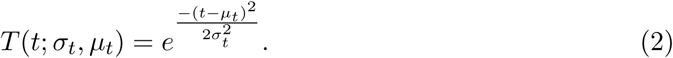

We refer to cells that were better fit by eq. 1 than by a constant term (just *a*_0_) as *time cells*, subject to several constraints described in detail below.

The mean of the time term *μ*_*t*_ was allowed to vary between −100 ms and 1700 ms and the standard deviation *σ*_*t*_ varied between 0 and 5 s. In order to ensure that *p*(*t*; Θ) can be considered as a probability we had to ensure that its values are bounded between 0 and 1. Therefore, the coefficients were bounded such that *a*_0_+*a*_1_ ≤ 1. The likelihood of the fit was defined as a product of these probabilities across all 1600 time bins within each trial and across all trials. We expressed the likelihood in terms of the negative log-likelihood (nLL), therefore instead of a product, a sum of the probabilities was computed:

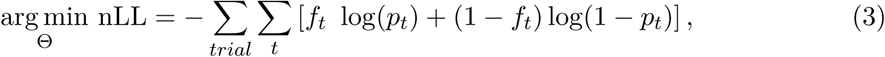

where *f*_*t*_ is the spike train.

In order to quantify whether the contribution of the terms that contained time was significant, the maximum log-likelihood was computed again, but this time with the time term set to zero (*a*_1_ = 0), such that the likelihood was affected only by the constant term *a*_0_. Since the models with and without time are nested, the likelihood-ratio test was used to assess the probability that adding the time term significantly improved the fit. The test is based on the ratio of the likelihoods of two different models and expresses how many times the data are more likely under one model than the other and it takes into account the difference in the number of parameters. To ensure that a unit will not be classified as a time cell only due to its activity in a single trial, the analysis was done separately on even and odd trials. For a unit to be classified as a time cell it was required that the likelihood-ratio test was significant (*p* < 0.01) for both even and odd trials. In order to eliminate units with ramping or decaying firing rate during a delay interval, *μ*_*t*_ was required to be within the delay interval and at least one *σ*_*t*_ away from either the beginning or the end of the interval. Also, to eliminate units with overly flat firing rate from classification as a time cell, *σ*_*t*_ was required to be at most equal to the length of the delay interval.

### Quantifying category specificity

The subset of units that passed the above criteria were classified as time cells. We then tested whether these units were also modulated by the category of stimulus on each trial (i.e., cats, dogs, sports cars, sedans) or for category sets (i.e., animals/cars). Category specificity was tested with a model that allowed four parameters, rather than one as above, to modulate the Gaussian-shaped time field, determined by the identity of the stimulus category on each trial. The probability of a spike at time point *t* was given as:

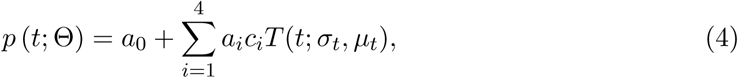

where *a*_0_ to *a*_4_ are the parameters to be estimated, while *μ*_*t*_ and *σ*_*t*_ are those estimated from the previous fit with a single time field eq. (1). The factor *c*_*i*_ was equal to 1 for trials when a stimulus from *i*-th category was presented and 0 otherwise. For instance, *c*_1_ = 1 for trials which started with a sample stimulus from the DOG category and *c*_2_ = 0 for trials where the sample stimulus was a CAT, SPORTS CAR or SEDAN). The model that includes category specificity eq. (4) and the model with a single time field eq. (1) are nested. Therefore, we use the likelihood-ratio test to assess the probability that adding the category specificity significantly improves the fit. When the outcome of the likelihood ratio test was significant (*p* < 0.01), a time cell was classified as category specific.

### Quantifying category set specificity

The category set specificity of the time cells was tested in analogous way to the category specificity, but using two time fields instead of four. Each of the two time fields corresponded to a particular category set. Thus the probability of a spike at time point *t* was given as:

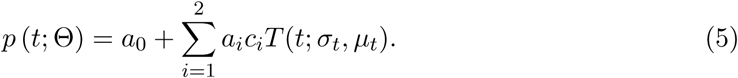

The factor *c*_*i*_ was equal to 1 for all the data points at trials when a stimulus from *i*-th category set was presented and 0 otherwise. For instance, *c*_1_ = 1 for trials which started with a sample stimulus from the DOG or CAT categories, and *c*_1_ = 0 for trials which started with a stimulus from the SPORTS CAR or SEDAN category. As in the case of the category specific cells, *μ*_*t*_ and *σ*_*t*_ were used as estimated from the fit with a single time field eq. (1). The likelihood-ratio test was used again to assess whether adding the category set specificity (eq. (5)) provided a better fit (*p* < 0.01) than the model with four time fields.

As an additional control to evaluate whether category set specificity was meaningful, we evaluated whether the number of category set specific time cells was more than one would expect from artificial pairings of categories. This was done by comparing the number of time cells that distinguished between animal and car category sets to the number of time cells that distinguished between artificial (not meaningful) mixtures of categories. One control model estimated the number of units that distinguished DOG and SPORTS CAR stimuli from CAT and SEDAN CAR stimuli. The other estimated the number of units that distinguished DOG and SEDAN CAR stimuli from CAT and SPORTS CAR stimuli. Because these artificial “category sets” are not meaningful, if the actual number of units coding for the true category sets—ANIMAL *vs* CAR—exceeds the number for the artificial category sets, we can conclude that the population contains information about the organization of the stimuli into category sets.

### Quantifying sustained activity

In order to analyze these results in the light of previous studies that argued for sustained stimulus-specific firing, we also identified units that distinguished category identity, but did not satisfy the criteria to be considered time cells. We found all the units that code for category identity by fitting a model in which the probability of a spike at time point *t* depended on a constant term that depended on the category identity of the sample stimulus:

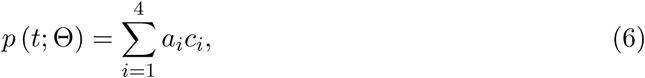

where *a*_1_ to *a*_4_ are the parameters to be estimated, and as above, *c*_*i*_ conveys category identity. Units were considered as category specific if a better fit was obtained with a model in eq. (6) than with a model that contained only one constant term *a*_0_. A subset of these units were also identified as category specific time cells using the analysis described above.

### LDA for cross-temporal classification

In addition to maximum likelihood approach, we used linear discriminant analysis (LDA) to quantify the decoding accuracy. We divided the 1.6 s interval composed of sample and delay period into 50 ms long non-overlapping time bins. We used the entire population composed of 500 units. The units were recorded during multiple recording sessions with different number of trials. To even the number of trials across all units we restricted the number of trials to the lowest number recorded from a single unit, 857 trials. For each time bin we trained an LDA classifier on 80% of randomly chose trials and used the remaining 20% of the trials for testing. The objective of classification was to accurately assign each trial to one of the four stimulus categories, with chance level being 25%. The testing was done on the same time bin as the training (to evaluate the decoding accuracy) but also on every other time bin (to evaluate performance of a classifier as a function of temporal distance between training and testing time bin). We repeated the training and testing for 10 iterations in order to obtain robust results (quantified through standard error of the mean). The classifier was implemented using Matlab 2017b function *classify*. To ensure stability of LDA the dimensionality of the training and testing data was reduced to full rank at before each run of the classifier.

### Computational model

The computational model used here is based on a previously-published method for computing a scale-invariant neural timeline (Shankar & Howard, 2012, 2013). The model can be understood as a two-layer feedforward network (Figure 1A). The first layer implements an approximation of the Laplace transform of the input to the network (keeping only the real part of the coefficients). The second layer approximates an inversion of the transform using the Post approximation formula (Post, 1930). Here we assume that the input function is a transient that captures information about the identity of the sample stimulus at the time it is presented. After the input is presented, the first layer codes the Laplace transform of the input function. As the delay progresses, this input function contains the identity of the sample stimulus further and further in the past. The Laplace transform contains this information and the second layer approximates a reconstruction of this function, with different units supporting different parts of the time axis. We compare the properties of units in the second layer of the network to experimentally-observed stimulus-specific time cells. These units estimate the time of the sample stimulus with decreasing accuracy as it recedes into the past, resulting in broader time fields and fewer units with time fields as the sample stimulus becomes more temporally remote.

**Figure 1.**
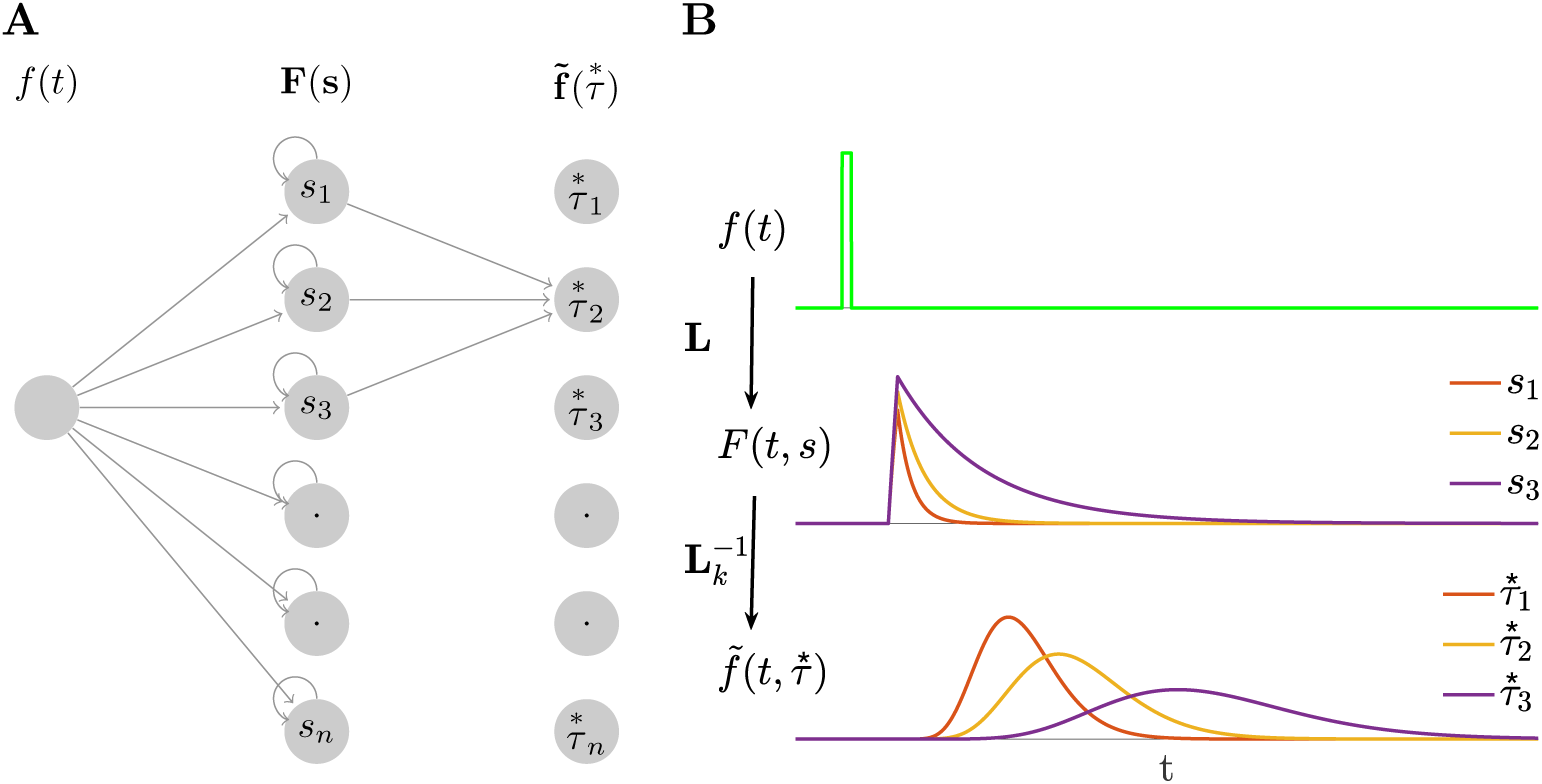
Constructing a scale-invariant compressed memory representation through an integral transform and its inverse. **A**. A schematic of the network architecture. The input stimulus *ƒ*(*t*) feeds into a layer of leaky integrators *F*(*t, s*) with a spectrum of time constants 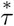 constituting a discrete approximation of an integral transform. *F*(*t, s*) projects onto 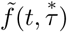 through a set of weights defined with the operator denoted as 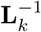 which implements an approximation of the inverse of the Laplace transform. Notice that the 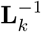 operator projects only a local neighborhood (*k* units) from each node in *F* layer to each node in 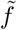 layer. **B**. A response of the network to a delta-function input. Only three nodes in *F*(*t,s*) are shown. Nodes in 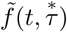 activate sequentially following the stimulus presentation creating a memory representation. The width of the activation of each node scales with the peak time determined by the corresponding 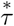, making the memory scale-invariant. Logarithmic spacing of the 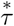 means that the memory representation is compressed.

To make this more concrete, in the present experiment with four distinct categories of stimuli, it is sufficient to consider an input vector **f**(*t*) consisting of 4 elements. At each moment **f**(*t*) gives the vector-valued category of the stimulus currently presented. When a stimulus from a particular category *A* is presented, the component *f*_*A*_ is set equal to 1 for a brief moment (i.e., a delta function input) and 0 at other times during the delay.

For simplicity, let us first consider the activity of units that receive input only from one component of **f**(*t*), say *f*_*A*_(*t*), with the understanding that in general units will receive some mixture of inputs from all four categories. Units in the first layer receiving input from *f*_*A*_(*t*), which we denote as *F*_*A*_(*t, s*), act as leaky integrators (first order low-pass filters):

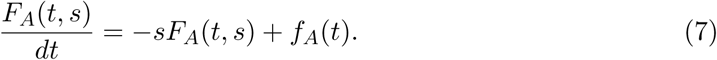

After receiving an input, units participating in *F*_*A*_ decay exponentially with a rate constant *s*(Figure 1B, middle). Each unit has a unique rate constant and we assume that the probability of observing a unit with rate constant *s* goes down like *1/s* (Shankar & Howard, 2013; Howard et al., 2015; Howard & Shankar, in press).

Let us denote the activity of units in the second layer receiving input from *F*_*A*_(*t*) as 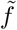_*A*_(*t*,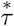), where 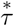 = — *k*/*s* and *k* is a positive integer that is common across all units. These units combine inputs from nearby values of *s* in *F*_*A*_, computing a *k*_th_ order derivative with respect to *s*:

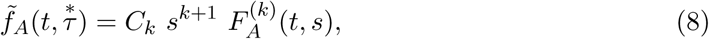

where *C*_*k*_ is a constant that depends only on *k*. Post (1930) proved that in the limit as *k* goes to infinity, 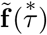 approximates 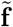.

When the input is a delta function at time zero, the activity of units in 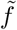_*A*_(*t*, 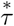) obey

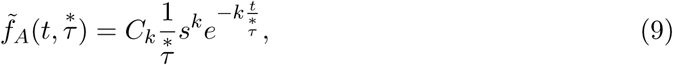

where *C*_*k*_ here is another constant that depends only on *k*. This expression is the product of an increasing power term 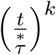 and a decreasing exponential term 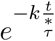. Consequently, the activity of each node in 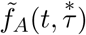 peaks at its value of 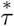 (Figure 1B, bottom).

It turns out (Shankar & Howard, 2012) that the width of each unit’s activity as a function of time depends linearly on its value of 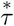 with a Weber fraction that is determined by the value of *k*. We found a good correspondence to the empirical data with *k* = 15. While *k* = 15 yielded the best correspondence with the data, all choices yield similar qualitative results. To implement the *k*^th^ order derivative with respect to *s, 2k* + 1 neurons from the first layer need to project to each neuron in the second layer with an on-center/off-surround connectivity pattern. If the neurons in the first layer are anatomically organized by their time constant, a spatially local neighborhood of 2*k* + 1 neurons from the first layer projects to each neuron in the second layer. A higher order derivative could also be computed by stacking up layers that implement lower order derivatives. For instance, a 6^th^ order derivative could be implemented by stacking up three layers each implementing a 2^nd^ order derivative. The values of 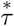 were logarithmically spaced between 100 ms and 1500 ms. Logarithmic spacing implements Weber-Fechner scaling.

To mimic stimulus-specificity we assumed that units in 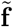 received a mixture of inputs from stimuli from four categories and two category sets (analogous to the behavioral task). For each unit, we picked one category as its preferred category and weighted its response by one (eq. 9) for that category. For the other categories, we picked coefficients randomly to weight the same temporal response. When a stimulus did not belong to the preferred category, but was from the same category set as the preferred category, we weighted the impulse response by a value taken from a normal distribution with mean 0.6 and standard deviation 0.3. When the stimulus was from the other category set, the temporal response was weighted by a value taken from a normal distribution with mean 0.3 and standard deviation 0.3. In addition, all of the coefficients were bounded between 0 and 1. These values were chosen informally to provide rough agreement with the empirical data.

## Results

In this study we analyzed recordings from 500 units in lateral prefrontal cortex (lPFC) of two macaque monkeys during performance of a delayed match to category working memory task initially reported in Cromer et al. (2010). The method is summarized in Figure 2. Neurons in lateral PFC (lPFC) are known to maintain stimulus information during the delay. Here we focus on the units that showed temporal modulation, with special attention to the existence of stimulus-specific time cells, as predicted by cognitive models of working memory (Howard et al., 2015). In order to facilitate comparison of the neural phenomena to theoretical models, we also include simulations of a computational model for a compressed neural timeline (Shankar & Howard, 2012, 2013; Howard et al., 2014).

**Figure 2.**
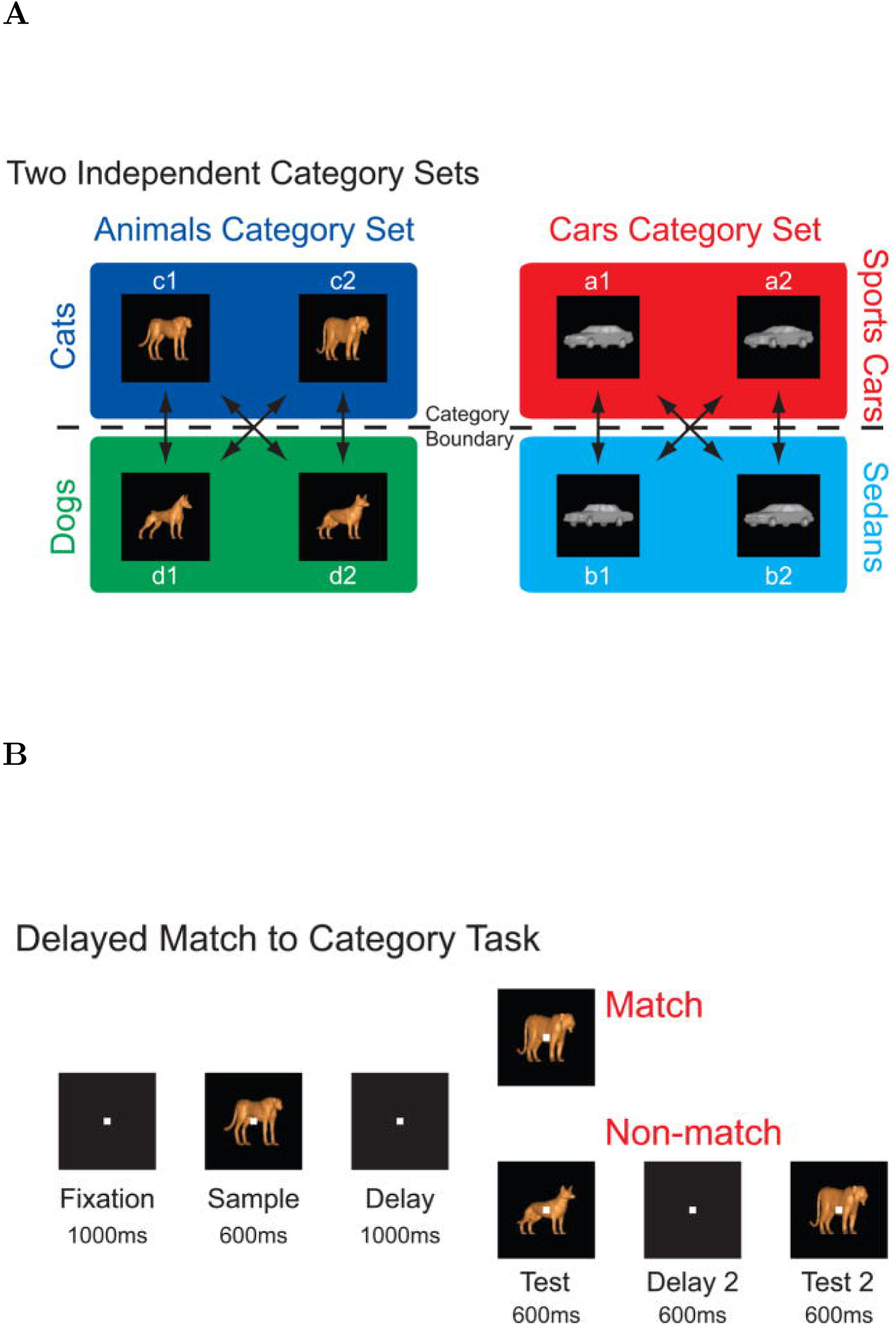
Behavioral Task. **A**. Stimuli were divided into two category sets, ANIMALS and CARS; each category set consisted of two categories. The ANIMAL category set consisted of CATS and DOGS and the CAR category set consisted of SPORTS CARS and SEDANS. Stimuli were morphed combinations of prototypes within a given category set. **B**. Two monkeys performed a delayed match to category task. On each trial, the monkey was required to respond to whether a test stimulus matched the category of the sample stimulus. To perform the task correctly the animal had to maintain a memory representation of the stimulus category throughout the sample and delay periods. Reproduced from Cromer et al. (2010).

### Units carrying temporal information

Out of 500 analyzed units, 240 were classified as time cells. Several examples of firing activity of those units are shown in Figure 3. The time cells activated sequentially, spanning the entire interval. The temporal profiles of all 240 time cells averaged across trials are shown on Figure 4A. The cells are sorted by the peak time of the estimated Gaussian shaped time fields (*μ*_*t*_).

**Figure 3.**
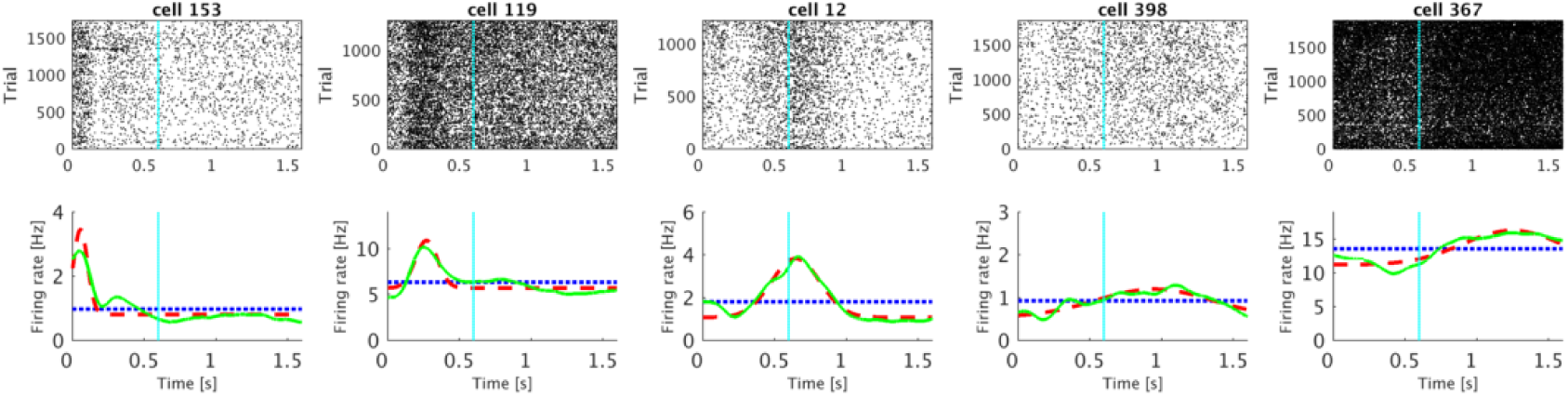
Representative examples of units classified as time cells. Each of the five columns shows activity of single unit. For each unit, the plot in the top row shows a raster of spikes across trials irrespective of the stimulus category. The bottom row shows the averaged trial activity (solid green line), the model fit with only a constant term (dotted blue line) and the model fit with a constant term and a Gaussian-shaped time field (dashed red line). On this and all following raster plots cyan line at 0.6 s marks the end of the sample and beginning of the delay period. See methods for details. The units were chosen such that the estimated peak time (*μ*_*t*_) increases progressively from the first unit to the fifth unit.

**Figure 4.**
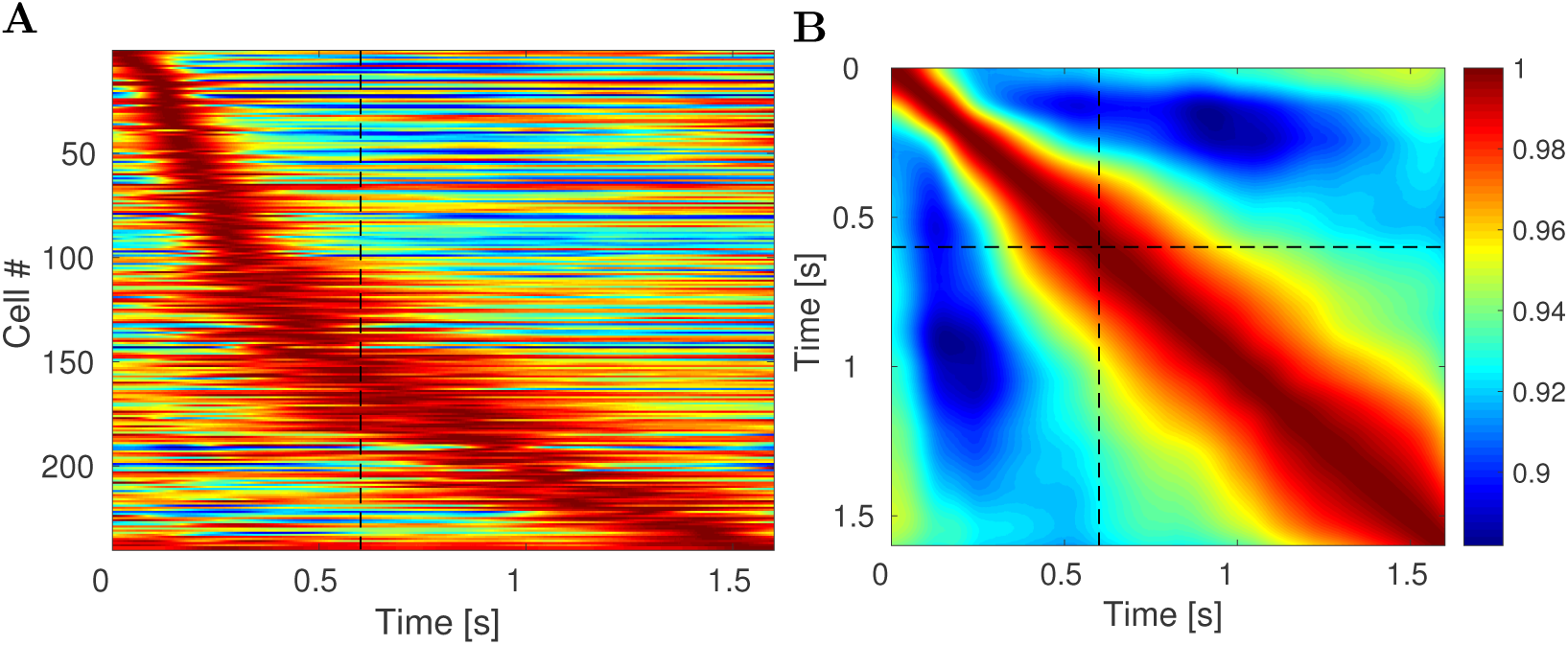
lPFC time fields show decreasing temporal accuracy for events further in the past. **A**. Activity of all 240 units classified as time cells during the 1.6 s interval. Each row on the heatplot corresponds to a single unit and displays the firing rate (normalized to 1) averaged across all trials. Red corresponds to high firing rate; blue corresponds to low firing rate. On this and all following heatmap plots dashed black line at 0.6 s marks the end of the sample and beginning of the delay period. The units are sorted with respect to the peak time estimated for their time field. There are two features related to temporal accuracy that can be seen from examination of this plot. First, time fields later in the delay are more broad than time fields earlier in the delay. This can be seen as the widening of the central ridge as the peak moves to the right. In addition the peak times of the time cells were not evenly distributed across the delay, with later time periods represented by fewer cells than early time periods. This can be seen in the curvature of the central ridge; a uniform distribution of time fields would manifest as a straight line. **B**. Ensemble similarity of all 240 time cells given through a cosine of the angle between normalized firing rate population vectors. The angle is computed at all pairs of time points during the observed interval. The bins along the diagonal are necessarily equal to 1 (warmest color). The similarity spreads out indicating that the representation changes more slowly later in the observed interval than it does earlier in the observed interval.

### Temporal information was coded with decreasing accuracy as the trial elapsed

There are two ways that a population of time cells would show decreasing temporal accuracy. First, the width of time fields should increase as the trial elapses. Second, the number of units with time fields earlier in the delay should be larger than the number later in the delay. Both of these properties were observed.

First, the width of the central ridge in Figure 4A increases from the left of the plot to the right of the plot, suggesting that units that fire earlier in the trial tend to have narrower time fields than the units that fire later. This impression was confirmed by analyses of the across-units relationship between the peak time (*μ*_*t*_) and the standard deviation (*σ*_*t*_) of the estimated Gaussian shaped time fields (Figure 5A). The correlation between the peak time and the width was significant (Pearson’s correlation 0.48, *p* < 10^−14^). A linear regression model linking the peak time (independent variable) and the width (dependent variable) gave an intercept of 0.09 ± 0.01 (mean ± SE), *p* < 10^−12^, and a slope of 0.15 ± 0.02, *p* < 10^−14^. To the extent the relationship is linear, it confirms a key quantitative prediction of a scale-invariant timeline; the dashed green line in Figure 5A is the prediction derived from the theoretical model (see methods for details).^1^

**Figure 5.**
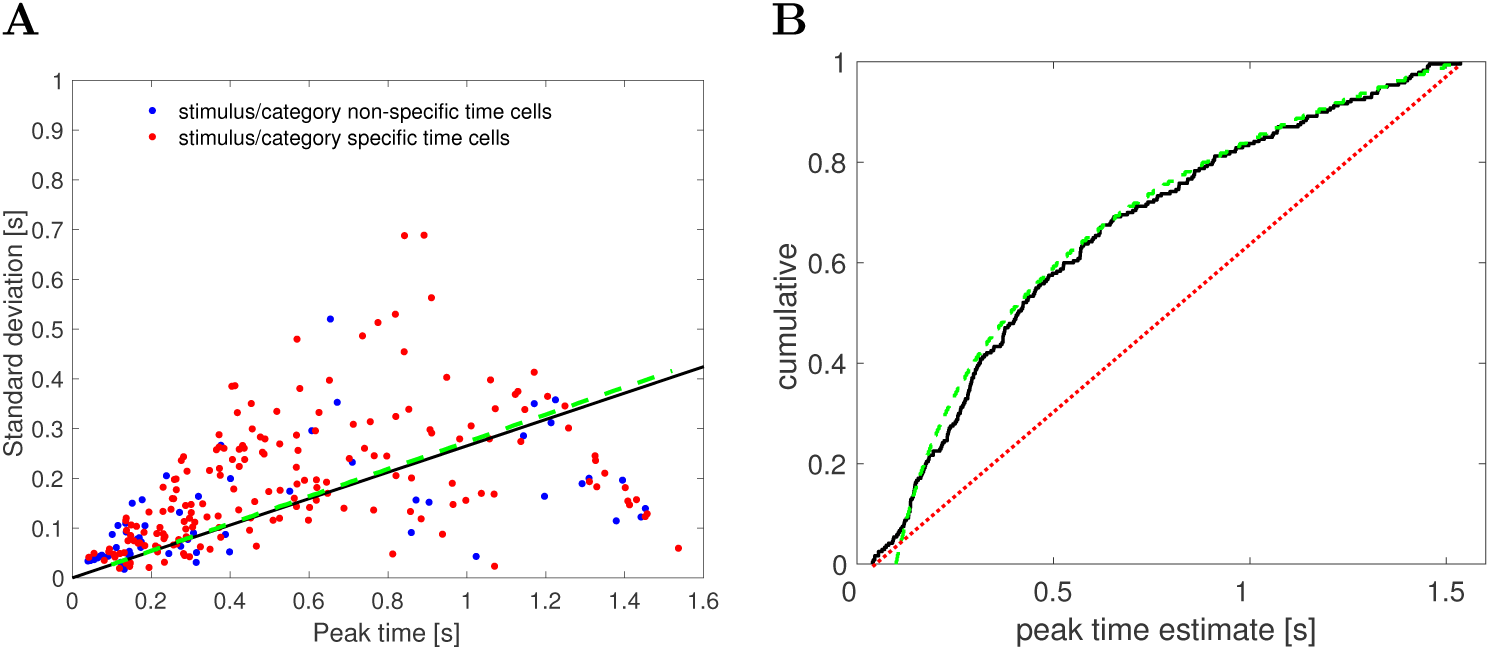
As the trial elapses, time cells in lPFC show broader and less frequent time fields. **A**. Width of the time fields increases with the peak time. Each dot represents the best-fitting parameters for a single unit classified as a time cell. There is no apparent difference between category/category set specific time cells (red) and the non-specific time cells (blue). The solid black line shows the results of linear regression (only the slope is shown, without the intercept term). The dashed green line is the relationship between the width and the peak time of time cells generated with the computational model with parameter *k* = 15. **B**. Peak times of the time fields are non-uniformly distributed along the delay interval. The cumulative density function for the parameter describing the peak firing of each time cell is shown as the solid black line. A fit with a uniform distribution is represented with a dotted red line. More time cells had time fields earlier in the delay interval and fewer had time fields later in the delay interval. The dashed green line shows the a power-law distribution with exponent −1 with values chosen between 100 ms and 1500 ms. The correspondence between the empirical results and this scale-invariant distribution is striking.

Second, the number of time fields later in the trial was smaller than the number of time fields earlier in the interval. This can be seen from the fact that the central ridge in Figure 4A does not follow a straight line, as would have been the case if it followed a uniform distribution. Rather, the curve flattens as the interval proceeds. To quantify this, we examined the distribution of the peak times across time cells (Figure 5B). The KS test rejected the hypothesis that the distribution of the peak times is uniform, *D*(240) = 0.28, *p* < 0.001. The dashed green line in Figure 5B is the cumulative that would be expected if the distribution was a power-law with exponent −1 with values between 100 ms and 1500 ms (choice of *k* does not affect this exponent). The correspondence of the observed results and this theoretical distribution suggests that the timeline is compressed logarithmically, consistent with the Weber-Fechner law. This prediction of the computational model is independent of the choice of *k*.

In addition, we investigated whether the data are better explained by fitting the stimulus presentation time (first 0.6 s of the observed interval) separately from the delay period (subsequent 1 s). We computed bilinear fit by finding two slopes and two intercepts that maximize the likelihood of the data given the bilinear fit. The power-law fit with the exponent of −1 explained the data better than the bilinear fit (since the two models had different number of parameters the fits were compared in terms of AIC and BIC: Δ*AIC* = 12.6, Δ*BIC* = 23).

The observation that temporal information was coded with decreasing accuracy as the trial elapsed was further supported by changes in the ensemble similarity across time. The ensemble similarity (Figure 4B) was computed for the population of time cells as a cosine of the normalized firing rate vectors between all pairs of time points during the observed 1.6 s long time interval.

### The population coding for time conjunctively carried information about category identity

Out of 240 units classified as time cells, 175 also distinguished the identity of the category of the to-be-remembered stimulus using the criteria described in the methods. Figure 6 shows several examples of such cells. These units were classified as time cells because they fired preferentially at circumscribed periods of time during the trial. However, the magnitude of their firing also depended dramatically on what category of stimulus was presented on the trial.

**Figure 6.**
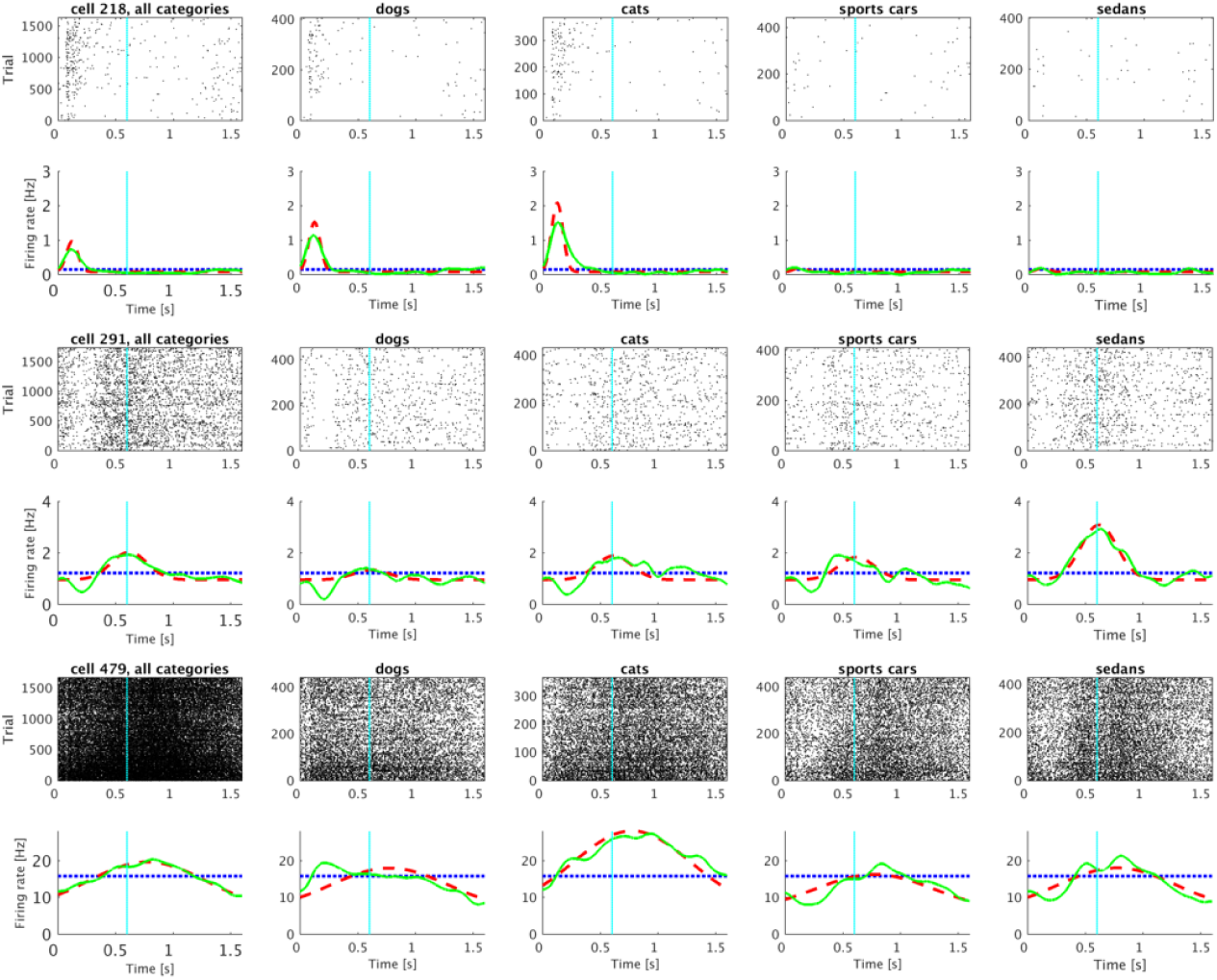
Representative time cells that were modulated by the category of the sample stimulus. A significant proportion of units classified as time cells distinguished the categories from one another. The activity of three category-specific time cells is shown (rows) with rasters corresponding to all the trials (left) as well as to each of the four stimulus categories (subsequent four columns). Category 1 and 2 are trials in which the sample stimulus was chosen from the dog or cat category respectively; Category 3 and 4 are sports car and sedan trials. Averaged trial activity is shown as a solid green line, the model fit with only a constant term is given by the dotted blue line and the best-fitting model—with different coefficients for each category—as a dashed red line.

### Time cells respected category-set structure

The two category sets (ANIMAL *vs* CAR) differ in their visual similarity. That is CAT and DOG stimuli are more visually similar to one another than they are to stimuli from the CAR category set. In order to determine whether stimulus-specific time cells respected this visual similarity structure, we noted that 45 out of 240 time cells met the definition for category-set specific time cells (see methods for details). Figure 7 shows several examples of representative units that were classified as category-set specific time cells.

**Figure 7.**
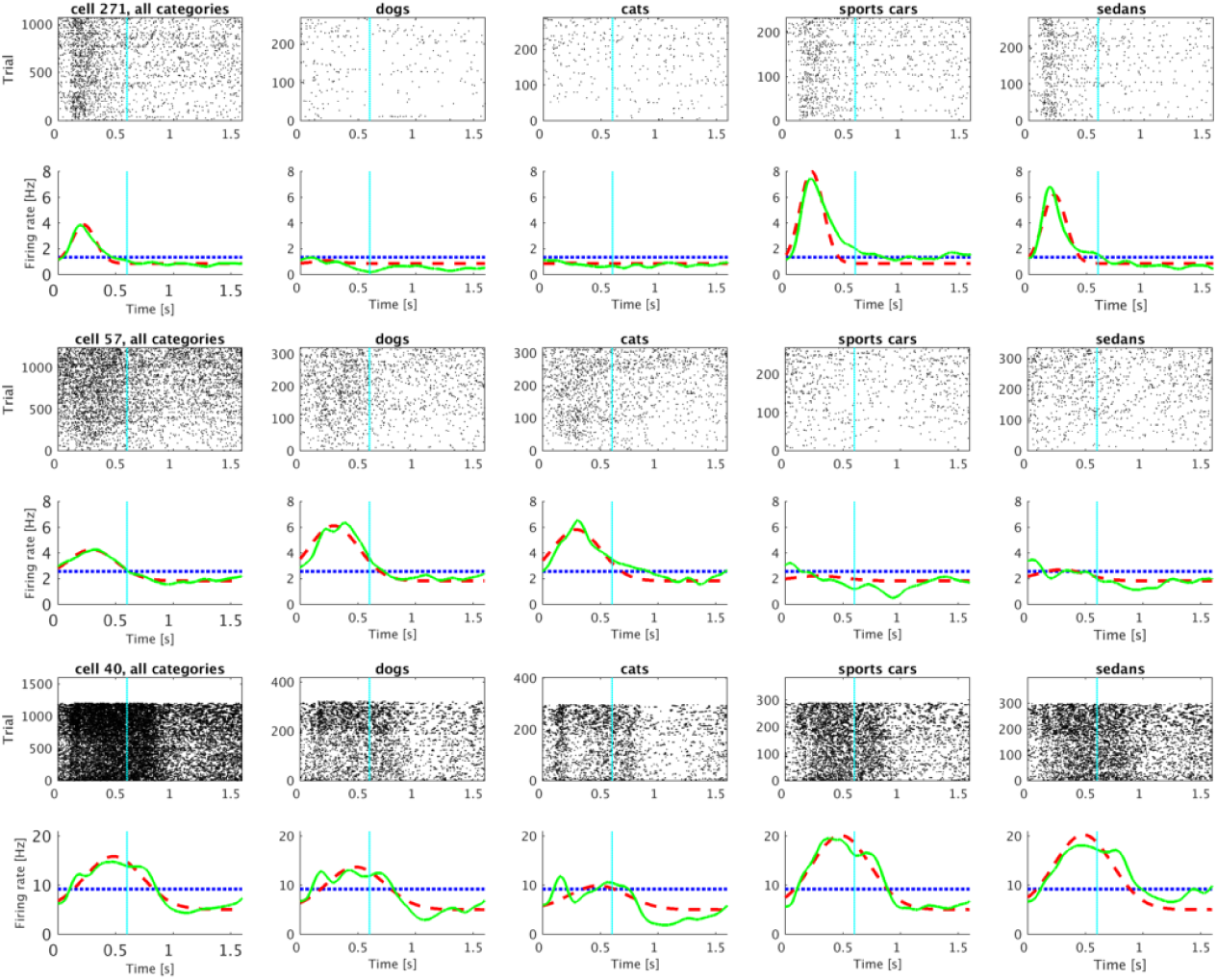
Representative time cells that were modulated by the category-set of the sample stimulus. As in Figure 6, each row is a cell. The left-most column shows data for all trials; the next four columns show data for trials in which the sample stimulus was chosen from each of the four categories. Category 1 and 2 correspond to the ANIMAL category set; Category 3 and 4 correspond to the CAR category set. The lines below each raster show averaged trial activity as a solid green line, the model fit with only a constant term as a dotted blue line and the best-fitting model—with distinct coefficients for each of the category sets—as a dashed red line.

As a control we computed the number of cells that had similar coefficients for artificial category sets. The number of category-set specific units significantly exceeded the number of cells specific for artificial category sets: 8 units distinguished DOG and SPORTS CAR stimuli from CAT and SEDAN stimuli. 15 units distinguished DOG and SEDAN stimuli from CAT and SPORTS CAR stimuli. Both of these proportions (8 and 15 out of 175) is reliably different than the 45/175 that distinguished the true category sets (ANIMAL vs CAR), *𝒳*^2^ = 28.81, *p* < 10^−7^ and *𝒳*^2^ = 16.91, *p* < 10^−4^.

The results for conjunctive coding of what and when information can be read off directly from the heatmaps in Figure 8, which shows the temporal profiles of all 175 stimulus specific time cells. The first heatmap Figure 8 shows the temporal profile for each unit in response to the category that caused the unit to fire at the highest rate. The middle heatmap shows the temporal profile for the same sorting of units for stimuli for the other category from the same category set. For instance, if a particular unit responded most to DOG stimuli, that temporal profile would be in the left heatmap and its response to CAT stimuli would be in the middle heatmap. The heatmap on the right shows the temporal profile for each unit in response to stimuli from the category set that did not include the unit’s best category. For instance, if a unit responded best to DOG stimuli, its profile in the right heatmap gives its response to stimuli from the CAR category set, including trials with sample stimuli from both the SPORTS CAR and SEDAN categories.

**Figure 8.**
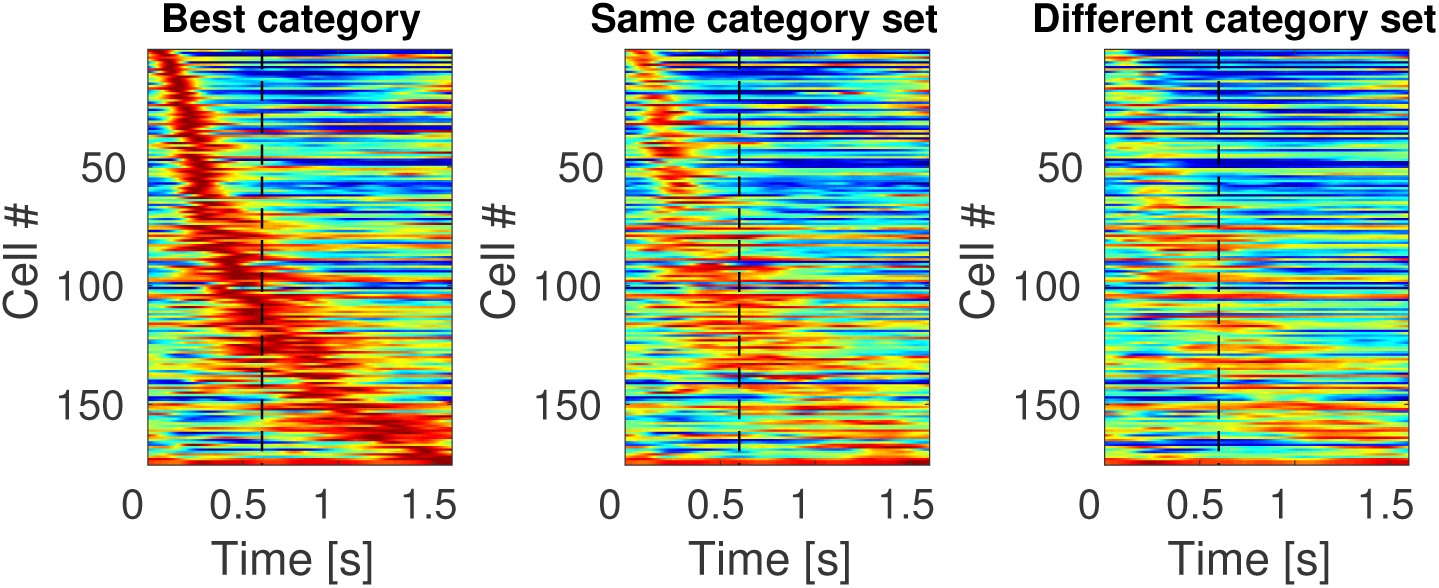
Sequentially activated time cells in lPFC encode time conjunctively with stimulus identity. The three heatmaps each show the response of every unit classified as a time cell. The heatmap on the left (“Best category”) shows the response of each unit to the category that caused the highest response for that unit, sorted according to the units estimated time of peak activity. The second column (“Same category set”) shows the heatmap for the same units, but for the other category from the same category set as that unit’s “Best category.” For instance, if a unit responded the most on trials in which the sample stimulus was chosen from the CAT category, then that units response to CAT trials would go in the first column and its response to DOG trials would go in the second column. The third column shows the response of each unit to trials on which the sample stimulus was from the other category set. Continuing with our example, a unit whose best category was CAT would have its response to CAR trials in the third column. The scale of the colormap is the same for all three plots and it is normalized for each unit such that red represents the unit’s highest average firing rate and blue represents its lowest average firing rate across time bins.

The sensitivity of the stimulus-specific time cells to category set can be noted from observing the difference between the heatmap in the second column and the heatmap in the third column. Although a difference between the first and second heatmaps could simply be a selection artifact, the difference between the second and the third indicates that time cells in this experiment respected the structure of the category sets.

### Majority of units that encoded category identity were time cells

The fit with four constant terms was better than the fit with just one constant term for 342 out of 500 units. These 342 units distinguished category identity. The majority of these units also showed temporally modulated firing. 269/342 of the category selective units were also temporally-modulated (Figure 9B). The firing dynamics of the remaining 73 category-specific units seemed irregular rather than sustained in time over the delay (Figure 9A). Figure 10 shows rasters for typical cells that were category specific but did not pass the threshold for reliable temporal modulation (as modeled by a Gaussian time field). Notice that the units that were fitted better with a Gaussian-shaped time field than with a constant term were not necessarily considered as time cells. This is because for classifying unit as a time cells we imposed an additional set of requirements regarding the peak time and standard deviation as described in the methods section (those criteria were necessary to fully define the time cells in terms of peak time and width of the temporal fields).

**Figure 9.**
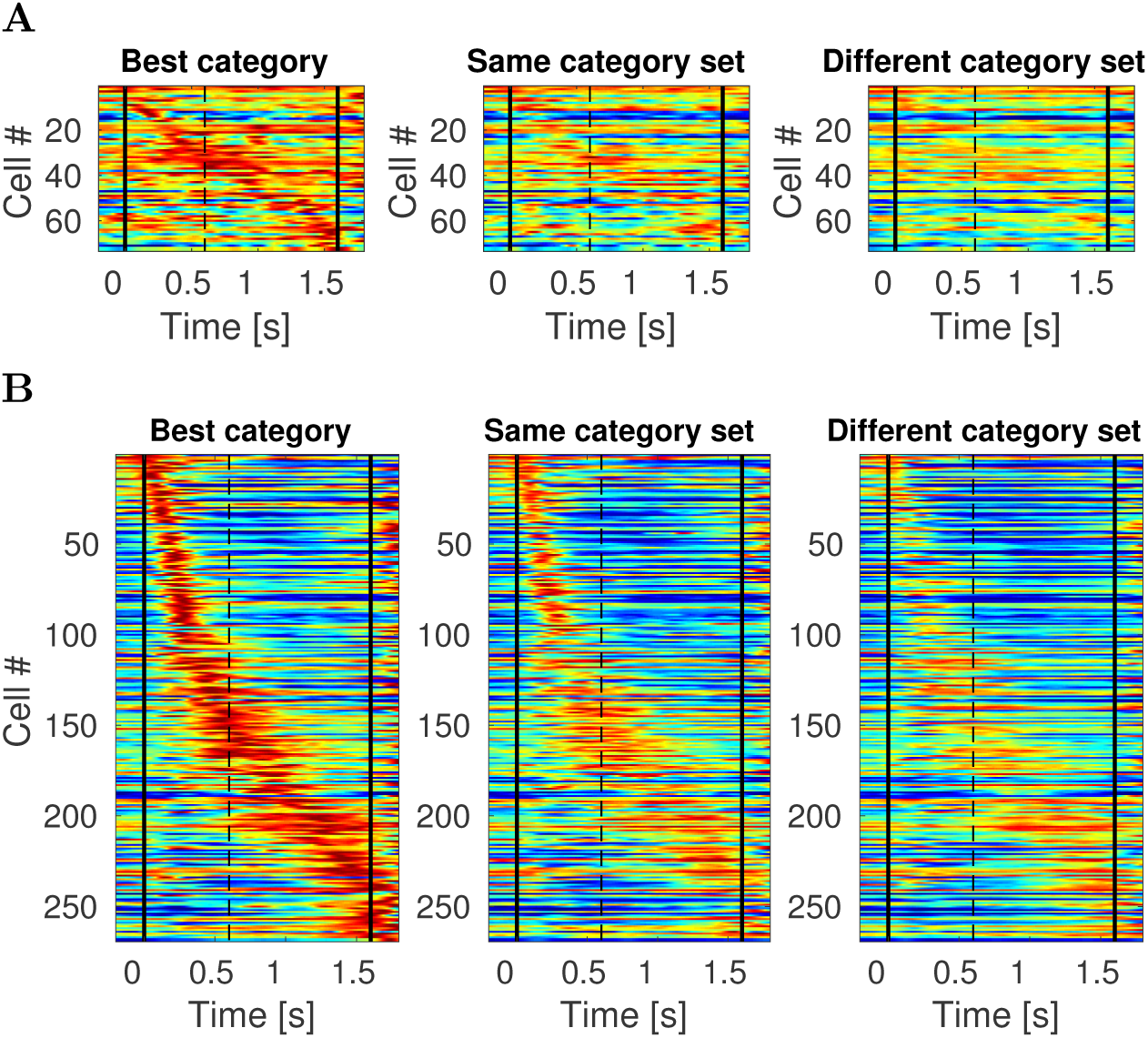
Category specific cells in lPFC exhibit temporally modulated firing more than stable persistent firing. Colormap and cell ordering is analogous to the one in Figure 8. Vertical black lines denote start of the stimulus presentation and end of the delay interval. **A**. Each of the three heatmaps shows activity of all 73 units that were category specific and where the fit with a constant term was better than the fit with a Gaussian-shaped time field. Most of these units are showing some form of temporally modulated firing, very few units are showing activity that could be considered as sustained throughout the entire stimulus presentation and delay interval. **B**. Activity of all 269 units that were category specific and fitted better with a Gaussian-shaped time field than with a constant term. This is a superset to the category specific time cells show in Figure 8, since to classify cells as a time cell we imposed an additional set of requirements. Some of these units that were not classified as time cells (93 of them) show ramping or decaying activity (which could mean that they would potentially be time cells if the delay interval was longer).

**Figure 10.**
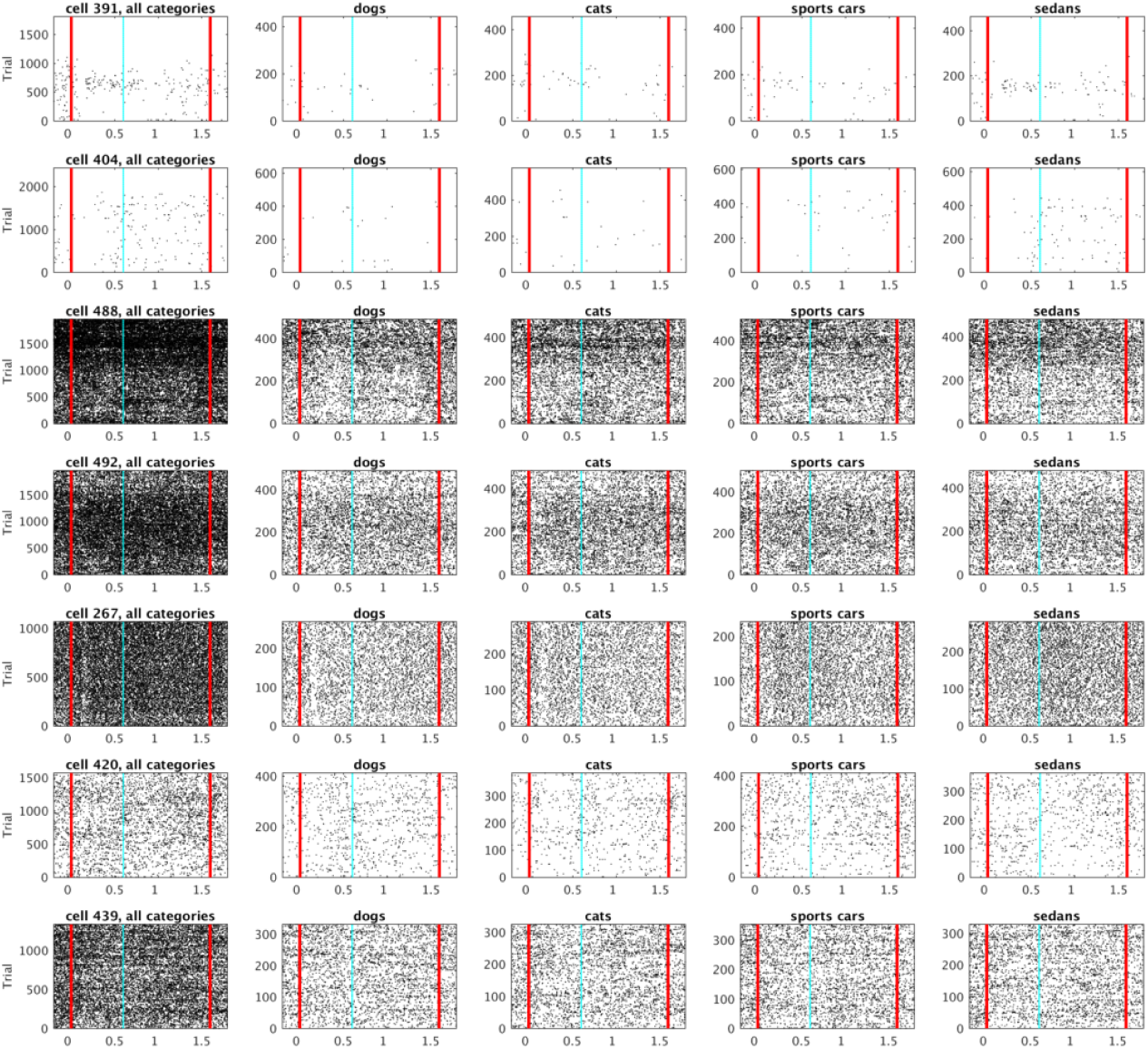
Examples of cells from figure Figure 9A that visually appear most similar to category specific sustained firing. Vertical red lines denote start of the stimulus presentation and end of the delay interval.

### The ensemble contained information about the category identity well above chance at almost all time points

Linear discriminant analysis (LDA) was performed to decode category identity of the sample stimulus at each 50 ms time bin of the sample and delay intervals (see methods). Accuracy for the majority of the time bins was above chance (Figure 12). The accuracy was computed for 10 runs, in each run trials were randomly assigned to train the model (80%, 686 trials) or held out for testing the classifier (20%, 171 trials). For the held out trials, if the classifier successfully classified 55 or more trials, this would exceed the Type I error rate at the .05 level. As a conservative estimate of decoding accuracy, we took time points for which the average over the 10 runs exceeded this value as reliably coding category identity. The accuracy was particularly high for the time bins where the classifier was tested on the same time bin it was train on (the diagonal in Figure 12). Every time bin after 100 ms along the diagonal was classified above chance (30/32 bins), indicating that the ensemble maintained information about category identity.

**Figure 11.**
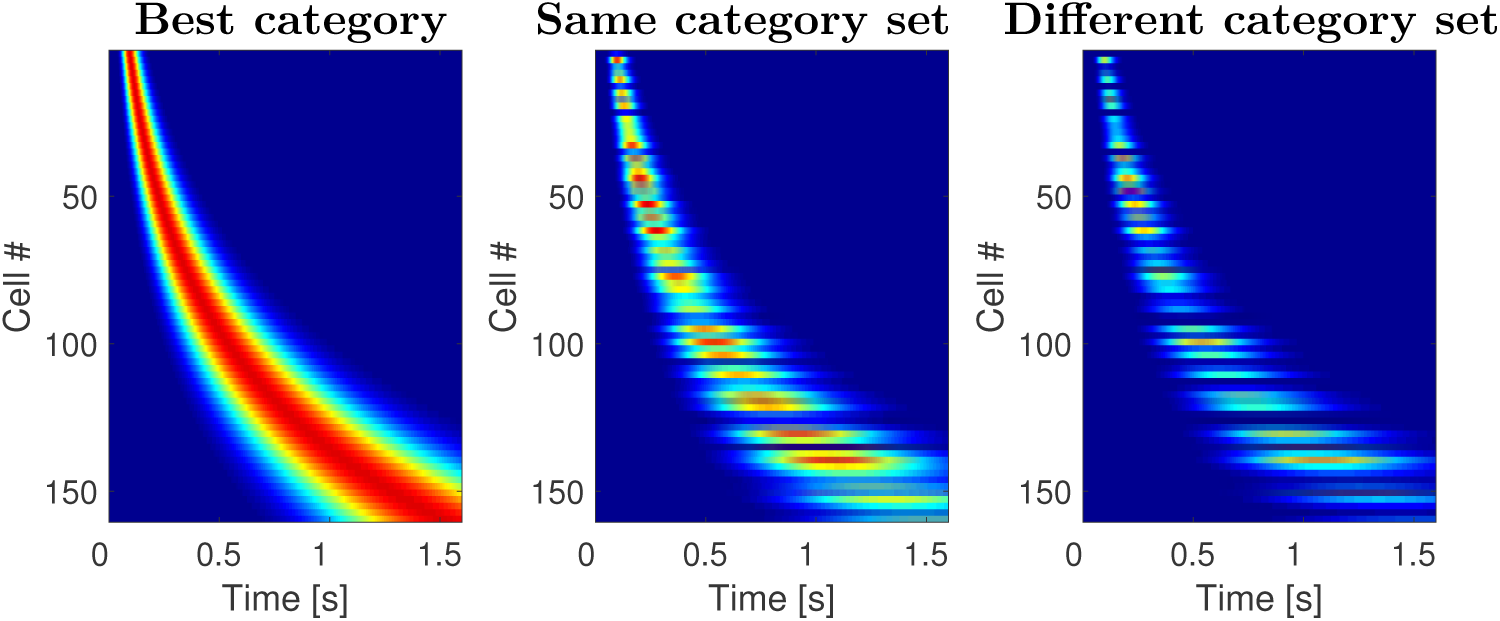
Sequentially activated time cells generated with the computational model. The three plots on the figure resemble the results shown on Figure 8. Analogous to the heatmaps in Figure 8, each row corresponds to a single model unit and displays its normalized activity across time. The cells are sorted with respect to the peak time. The two features observed in Figure 8 are fully captured by the model: the time fields later in the delay were more broad than the time fields earlier in the delay and the density of time fields decreased as a function of time. This illustrates that the model can indeed account for the firing dynamics of the sequentially activated time cells that form a compressed representation of time. In addition, the model predicts that the stimulus selectivity observed in the data. This is because every time a stimulus is presented, it activates not only its own memory representation, but also the memory representation of other stimuli to the degree they are similar to the presented stimulus. The response to stimuli from the same category set is on average more similar the response to stimuli from different category sets.

**Figure 12.**
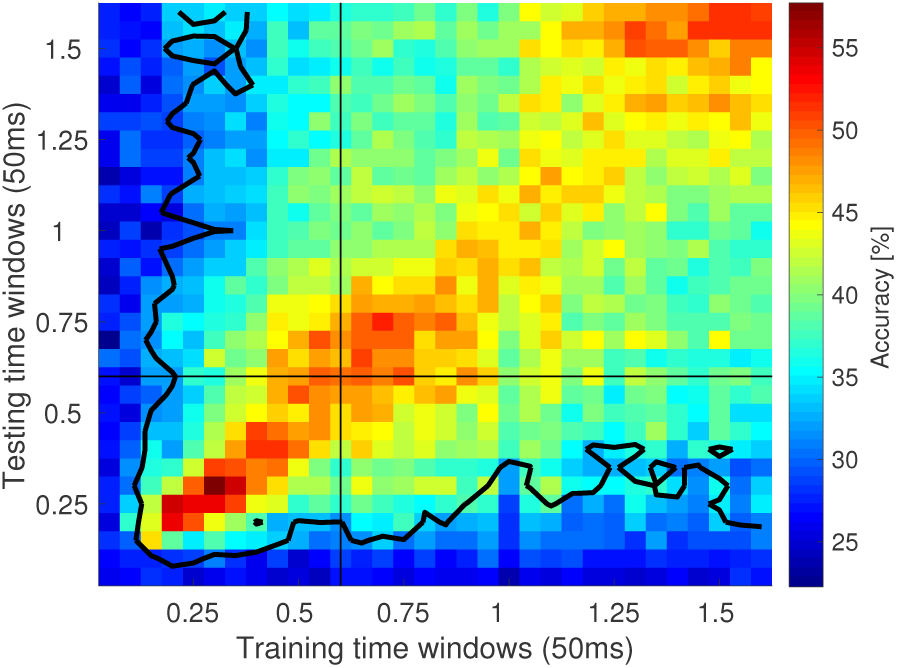
Decoding category identity using linear discriminant analysis (LDA) reveals gradual change of the population code across time. The heatmap displays the accuracy of the LDA classified applied on 50 ms time bins. Each bin provides classification accuracy for the classifier trained on a bin denoted on x-axis and tested on a bin denoted on y-axis. Each bin is computed by averaging across 10 runs in which training and test trials were randomly chosen (80% of trials were used for training with 20% of trials held out for testing). The contour encloses bins where classification accuracy averaged over the 10 runs exceeded the Type I error rate at the .05 level.

The performance of LDA decoder appeared decreased as a function of temporal distance between the time bin the decoder was trained on and the time bin the decoder was tested on. This is indicated by the gradual change in columns of the heatmap in Figure 12. Peak accuracy was obtained around the diagonal elements and gradually decreased for points further from the diagonal, suggesting that the part of the neural ensemble decoding stimulus category changed gradually over the delay. This observation is consistent with the gradual change through sequential activation observed in time cells.

### Computational model for compressed memory representation

Previous work in computational neuroscience (Shankar & Howard, 2012; Howard et al., 2014) and cognitive psychology (Shankar & Howard, 2012; Howard et al., 2015) has developed a quantitative model for how a compressed timeline could be constructed. This method (described in more detail in the methods) makes a strong commitment to scale-invariance of the temporal representation inspired by robust behavioral results from timing and memory experiments (Rakitin et al., 1998; Howard, Youker, & Venkatadass, 2008). Consistent with these behavioral findings, and the logarithmic compression of receptive fields in the visual system (Schwartz, 1977; Van Essen, Newsome, & Maunsell, 1984), theoretical considerations (Shankar & Howard, 2013; Howard & Shankar, in press) strongly suggest that time fields should also be logarithmically compressed. This quantitative argument makes two clear predictions. First, the width of time fields should go up linearly with the time of their peak. Second, the number of time fields centered on a time *τ* should go down like *τ*^−1^.

This model can be understood as a two-layer feedforward network. However, because it can be well-described mathematically, analytic results can be readily obtained (see methods for details). Figure 5 provides a means to evaluate whether these quantitative predictions are consistent with the empirical data. The straight line relating spread of the time fields to their center (dashed green line in Figure 5A) is a qualitative prediction of the model; the slope of the line is controlled by a single parameter *k* (see methods for details). The distribution of time fields given by the model is given by the distribution *N*(*τ*) ≃ *τ*^−1^; the dashed green line in Figure 5B shows this distribution with two parameters controlling the smallest and largest possible values of the center of the time field (set to 100 ms and 1500 ms respectively). The agreement of the empirical data with the predictions of a scale-invariant timeline is very strong.

In order to further evaluate the comparison between the model and the empirical results, we also generated heatmaps as in Figure 8. By causing different units in the timeline to respond to differentially to stimuli from different categories, we were able to generate a strong agreement with the empirical results, as shown in Figure 11. Because these are analytic results, there is no noise in the time fields across trials. This qualitative fit of the computational model supports quantitative findings from the parameters of the descriptive model of time cells and the linear classifier (Figure 12).

## Discussion

During performance of a working memory task, some neurons in lPFC fired in sequences while information needed to be maintained in working memory. The sequences exhibited coding of temporal information—the time since the sample stimulus was presented (Figure 4)—conjunctively with information about the identity of the category of the sample stimulus (Figure 8). The temporal information decreased in accuracy as time elapsed (Figure 5). This decrease in accuracy aligned well with predictions of a scale-invariant timeline taken from cognitive models of memory.

### Temporal information throughout the brain

The present findings provide robust evidence of conjunctive coding of what and when information in the lPFC. Although to our knowledge this is the first report of robust stimulus-specific time cells, many recent papers have shown evidence for sequentially activated time cells in a broad range of brain regions and multiple species. Previous studies in rodents have found time cells with similar properties in hippocampus (MacDonald et al., 2011; Kraus, Robinson, White, Eichenbaum, & Hasselmo, 2013; Salz et al., 2016; Terada, Sakurai, Nakahara, & Fujisawa, 2017), medial prefrontal cortex (Tiganj et al., 2016; Bolkan et al., 2017), and striatum (Akhlaghpour et al., 2016; Mello et al., 2015). A monkey study has previously observed sequentially-activated time cells in dlPFC and striatum (Jin et al., 2009). In some of those previous studies showing time cell activity, the experimental procedure would not enable measurement of conjunctive what and when information (Jin et al., 2009; Tiganj et al., 2016; Mello et al., 2015; Kraus et al., 2013). In some other studies it would have been possible in principle to measure robust conjunctive what and when information (Akhlaghpour et al., 2016), but it was not observed or reported. MacDonald et al. (2011) observed some evidence for conjunctive what and when coding in the rodent hippocampus, but it was not as reliable as the present study (see also MacDonald et al. (2013)). Terada et al. (2017) showed reliable evidence for stimulus coding in the rodent hippocampus. Systematic study will be necessary to determine under what circumstances time cells also show evidence for stimulus identity.

The presence of sequentially-activated time cells, regardless of whether or not they also code for stimulus identity, in so many brain regions with such similar functional properties is striking. This may point to a fundamental role for a compressed timeline in many different forms of memory. Our conventional understanding of the cognitive neuroscience of memory describes memory as composed of a number of separable systems associated predominantly with various brain regions (Squire, 2004; Eichenbaum, 2012; Jenkins & Ranganath, 2016). Although there are ongoing disputes about how exactly to specify the systems, there is broad consensus that the hippocampus is associated with the declarative memory system, the striatum is associated with a non-declarative implicit memory system and the PFC is associated with a working memory system, etc. The fact that such similar temporal representations are observed in regions associated with so many distinct memory systems suggests that these memory systems rely on a common form of temporal representation. Perhaps different memory systems perform different operations on a common representation (Howard et al., 2015; Aronov, Nevers, & Tank, 2017).

For many years, the default understanding of the neural basis of working memory was that working memory is maintained through stable persistent firing (Fuster & Alexander, 1971; Funahashi, Bruce, & Goldman-Rakic, 1989; Goldman-Rakic, 1995; Curtis & D’Esposito, 2003). We did not observe large numbers of category-selective cells that exhibited sustained firing (see Figure 10 for the best examples). In contrast, category-selective units were temporally modulated, either as time cells or as ramping or decaying cells. Thus this study adds to a growing body of empirical work (e.g., Spaak, Watanabe, Funahashi, & Stokes, 2017; Lundqvist et al., 2016; Stokes et al., 2013; Murray et al., 2016) that requires an updated view of working memory as a dynamically-changing representation.

Interestingly, even though this task did not test animals’ memory for temporal order, our results clearly show that the stimulus selective neurons nonetheless conveyed temporal information. Thus, maintaining a memory representation of what happened when over the recent past might be largely spontaneous.

We observed that the width of the temporal receptive fields increased soon after the stimulus onset, despite the fact that the stimuli remained present for 600 ms. Similarly, when LDA was used for decoding the stimulus identity, we observed the peak performance soon after the stimulus onset. This suggests that the stimulus onset is perceived as a more salient effect than the stimulus offset. Further research with experimental paradigms that include longer stimulus presentation is needed to evaluate how the stimulus duration is encoded. Furthermore, further research is needed to understand whether the neural timeline can maintain multiple stimuli simultaneously as well as multiple repetitions of the same stimulus. The theoretical framework described here is linear and predicts that multiple repetitions of the same stimulus result in additive response that allows reconstruction of the presentation time of each repetition.

### Computational model of working memory

Computational neuroscience studies of working memory have predominantly focused on maintaining information about the identity of presented stimuli, either through sustained (Goldman-Rakic, 1995; Curtis & D’Esposito, 2003; Wang, 2001) or through time-varying firing dynamics (Durstewitz & Seamans, 2006; Stokes, 2015; Lundqvist et al., 2011; Murray et al., 2016; Sreekumar, Dennis, Doxas, Zhuang, & Belkin, 2014). Even though very useful, such models do not readily account for the temporal aspect of memory. Other computational models (e.g., Goldman, 2009; Grossberg & Merrill, 1992; Itskov, Curto, Pastalkova, & Buzsáki, 2011) can also account for sequentially-active firing that can be used to read off temporal information.

The computational model used here differs from previous work in that it makes a strong commitment to a logarithmically-compressed timeline and is mathematically tractable, enabling straightforward derivation of behavioral predictions (Shankar & Howard, 2012, 2013). The biological plausibility of the key components of the model has been studied closely. The major objection, that the method requires neurons with slow functional time constants, has been addressed by showing that a single-cell model based on known properties of persistently-firing neurons (Egorov et al., 2002; Fransén, Tahvildari, Egorov, Hasselmo, & Alonso, 2006) can be readily adapted to generate a broad range of slow functional time constants (Tiganj, Hasselmo, & Howard, 2015; Tiganj, Shankar, & Howard, 2013). The other major assumption of the model is a feedforward projection that is functionally equivalent to a set of on-center/off-surround receptive fields placed in series. The logarithmic compression of temporal receptive fields parallels the logarithmic compression of visual receptive fields, which has been known for decades (Hubel & Wiesel, 1974; Van Essen et al., 1984). The observed compression is consistent with Weber-Fechner law, providing a potential neural substrate for this widely observed psychophysical law. Moreover the mathematics of this model can be readily adapted to provide a description not only of time cells, but a variety of findings from the place cell literature and even sustained firing (Howard et al., 2014).

### Stimulus-specific time cells are predicted by many theories of memory

Previous studies have reported that it is possible to extract temporal or stimulus identity information by applying different decoding techniques on the activity of neural populations (Stokes et al., 2013; Kim, Ghim, Lee, & Jung, 2013; Pesaran, Pezaris, Sahani, Mitra, & Andersen, 2002; Baeg et al., 2003; Hung, Kreiman, Poggio, & DiCarlo, 2005). Neural models of timing rely on gradually changing firing rate throughout a delay period rather than temporal receptive fields (Kim et al., 2013; Gavornik & Shouval, 2011; Simen, Balci, de Souza, Cohen, & Holmes, 2011). Similarly, recurrent neural networks, including liquid state machines (Buonomano & Merzenich, 1995; Maass, Natschläger, & Markram, 2002; Buonomano & Maass, 2009; White, Lee, & Sompolinsky, 2004) can be shown to maintain information about preceding stimuli, but the decoder necessary to extract that information into a useful form can be quite complex.

Fusi, Miller, and Rigotti (2016) have argued that the brain expends significant resources representing features conjunctively using mixed selectivity to enable linear decoding (Rigotti et al., 2013). For coding of continuous dimensions, mixed selectivity manifests as receptive fields. To make a concrete example, the hippocampal place code might have consisted of as few as two neurons that fire proportionally to the *x* or *y* position of the animal. Although this code would require few neurons to represent position, learning an association between a location in the middle of an arena and reward would be a significant computational challenge. Instead, the hippocampal place code uses many neurons, each with a place field; the set of all place fields tile the enclosure. Even though this coding scheme uses many more neurons it is computationally straightforward to learn an association between a circumscribed spatial position (represented by the currently-active place cells) and some behaviorally relevant outcome. Despite the fact that sequentially activated stimulus-specific time cells seem to require a great many neurons, conjunctive coding of what and when information (Figure 8) enables direct readout of the elapsed time and the stimulus identity.

## Acknowledgments

The authors gratefully acknowledge support from ONR MURI N00014-16-1-2832, NIBIB R01EB022864, NIMH R01MH112169, and NIMH R37MH087027.

## Footnotes

1 Other values of *k* would have also resulted in a straight line, but with a different slope. Smaller values of *k* result in a steeper slope.

